# Identification of QTL for leaf angle at canopy-wide levels in maize

**DOI:** 10.1101/499665

**Authors:** Dengguo Tang, Zhengjie Chen, Jixing Ni, Qin Jiang, Peng Li, Le Wang, Jinhong Zhou, Chenyang Li, Jian Liu

## Abstract

Leaf angle (LA) is one of the most important canopy architecture related traits of maize (Zea mays L.). Currently, there is an urgent need to elucidate the genetic mechanism of LA at canopy-wide levels for optimizing dense-planting canopy architecture. In present study, one RIL population derived from two parent lines which show distinct plant architecture was used to perform QTL mapping for LA at eight leaves below the tassel under three environments. Dozens of QTL for LA at eight leaves were identified, which were mapped on all maize chromosomes except for the tenth chromosome. Among them, there were nine common QTL as they were identified for LA more than 1 leaves or in two or three environments. And individual QTL could explain 1.29% - 20.14% of the phenotypic variation and affect LA of 1-8 leaves, including qLA5.1 affected LA of all eight leaves, qLA3.1 affected LA of the upper leaves (1stLA to 4thLA), and qLA9.1 could affect LA of the lower leaves (5thLA to 8thLA). Furthermore, the results indicated that the genetic architecture of LA at eight leaves was different. Specifically, 8thLA was mainly affected by major and minor QTL; 1stLA, 4thLA and 5thLA were affected by epistatic interactions beside major and minor QTL; while the other four LAs were simultaneously affected by major QTL, minor QTL, epistatic interactions and environments. These results provide a comprehensive understanding of genetic basis of LA at canopy-wide levels, which will be beneficial to design ideal plant architecture under dense planting in maize.

**Author contribution statement:** J. L. and D. T. designed and supervised the study, D. T., Z.C., J.N., Q.J., P.L., L.W., J.Z., C.L. performed the phenotypic data collection. D. T. analyzed the data and drafted the manuscript, D. T. and Z.C. revised and finalized the manuscript. All the authors read and approved the manuscript.

**Key message:** Dozens of QTL for leaf angle of eight consecutive leaves were identified in the RIL population across three environments, providing the information that optimization of canopy architecture at various canopy levels.

## Introduction

Maize (*Zea mays* L.) is one of the most important cereal crops worldwide. The primary goal of maize breeding programs is to generate high-yielding varieties. During the past several decades, an increase in maize yields was largely due to an increase in the plant density, rather than improvement of the potential yield per plant (Duvick, 2005; Ma et al., 2014b; Mock and Pearce, 1975; Russell, 1991; Tollenaar and Wu, 1999). In order to adapt to the high dense planting, a number of dramatic changes in plant architecture have been observed. Moreover, several key parameters 3 that affected the optimal plant architecture were determined, including upright leaves, maximum photosynthetic efficiency, and small tassel size (Mock and Pearce, 1975). Leaf angle (LA) is a critical parameter of plant architecture by impacting light interception and photosynthesis. Maize breeding practices had also shown that LA was an essential agronomic trait in the development and adoption of high-yielding varieties of maize. As the breeders focused on improving the grain yield, LA score has decreased remarkably, thereby reshaping the plant architecture from expanded to compact (Anderson and Denmead, 1969; de Wit, 1965; Duncan et al., 1967; Ku et al., 2010a). Comprehensive analysis of the correlations between the LA trait and grain yield revealed two interesting phenomenon: (1) although the LA had significantly decreased over the past several decades, smaller LA does not guarantee higher yield; and (2) further increase in light interception efficiency requires variable LA at various parts of maize plant (Duncan, 1971; Lambert and Johnson, 1978; Ma et al., 2014a; Mickelson et al., 2002; Pepper et al., 1977; Winter and Ohlrogge, 1973; Zhang et al., 2017). Recently, Mantilla et al. (2017) proposed that optimization of canopy architecture can be manipulated by varying LA at different canopy levels to achieve maximum production potential in cereal species.

Both QTL mapping and genome-wide association analysis (GWAS) were used to dissect the genetic basis of LA in maize; hundreds of quantitative trait loci (QTL) for LA have been identified throughout all ten maize chromosomes. These studies have significant variability in the numbers and node positions of selected leaves, statistical methods of phenotype characterization, types of mapping populations, and QTL mapping strategies. Detailed analysis of the previous studies indicated that various research groups selected different numbers and node positions of the leaves for QTL analysis. In most cases, three continuous leaves, including the ear leaf and the leaves above and below the ear, were selected for QTL analysis (Ding et al., 2015; Ku et al., 2016; Ku et al., 2012; Ku et al., 2010b; Li et al., 2015; Mickelson et al., 2002; Ming et al., 2007; Shi et al., 2017; Zhang et al., 2017); while, in some instances, the first leaf below the flag (Pan et al., 2017; Tian et al., 2011; Wang et al., 2017a; Yang et al., 2015b) or two leaves near the ear (Chen et al., 2015; Hou et al., 2015) were chosen. In addition, two statistical methods for phenotypic data were used in QTL mapping, one was that the average values of the leaves, the other was that the value of the individual leaf. Furthermore, different mapping populations were adopted, such as F_2:3_ (Chen et al., 2015; Hou et al., 2015; Ku et al., 2012; Ku et al., 2010b; Ming et al., 2007; Yu et al., 2006), F_4_ (Chen et al., 2015), RIL (Ku et al., 2016; Li et al., 2015; Mickelson et al., 2002; Shi et al., 2017; Wang et al., 2017a; Yang et al., 2015b; Zhang et al., 2017), Four-Way Cross Mapping Population (Ding et al., 2015), NAM (Tian et al., 2011) and ROAM (Pan et al., 2017). In combination with the QTL mapping, Tian et al. (2011) and Pan et al. (2017) identified 203 and 10 single-nucleotide polymorphisms (SNPs), respectively, associated with LA in the GWAS studies.

Identification of actual genes responsible for LA QTL and isolation of the mutants with altered LA is the critical step to unravel the genetic and molecular mechanisms underlying maize LA. To date, four genes *ZmTAC1* (Ku et al., 2011), *ZmCLA4* (Zhang et al., 2014), *ZmRAVL1* and *Zmbrd1* (Tian et al., 2019) located in the QTL regions for LA and six LA mutants, including *liguleless1* (*lg1*) (Moreno *et al*., 1997), *lg2* (Walsh *et al*., 1998), *Liguleless3-O* (*Lg3-O*) (Muehlbauer *et al*., 1999), *Liguleless narrow-Reference* (*Lgn-R*) (Moon *et al*., 2013), *droopingleaf1* (*drl1*), *and drl2* (Strable *et al*., 2017), have been cloned. *Lg1*, *lg2* and *lgn-R* mutants exhibit a defect in the ligule and auricle tissues and a decrease in leaf angle (Harper and Freeling, 1996; Moon *et al*., 2013; Sylvester *et al*., 1990; Walsh *et al*., 1998). Notably, *LG1*, *LG2* and *LGN* were shown to act in a common pathway involved in ligule development (Harper and Freeling, 1996; Moon *et al*., 2013). Similarly, the *Liguleless3-O (Lg3-O)* mutant also developed a decreased leaf angle, which may be due to a defect in the blade-to-sheath transformation at the midrib region of the leaf (Fowler *et al*., 1996; Muehlbauer *et al*., 1997; Muehlbauer *et al*., 1999). Nevertheless, the LAs in the *drl1* and *drl2* mutants are increased; the *drl* gene is required for proper development of the leaf, leaf support tissues, and for restricting auricle expansion at the midrib (Strable *et al*., 2017). More recently, Tian et al. (2019) had cloned two LA QTL, *UPA1* and *UPA2* (Tian et al., 2019); *UPA2*, which is located 9.5 kilobases upstream of *ZmRAVL1*, regulates expression of ZmRAVL1 as a distant cis-regulatory element; *UPA1* encode *a brassinosteroid C-6 oxidase* (*brd1*) gene, which is participating in the synthesis of brassinosteroid. The authors proposed a leaf angle regulating model which composed of *UPA2, UPA1* and brassinosteroid and verified that by manipulating *ZmRAVL1* or using favorable alleles in wild relatives can generate upright leaf architecture and further high yield hybrids under dense planting (Tian et al., 2019).

In this study, one RIL population developed in the previous study was used for QTL mapping of LA of eight consecutive leaves below the tassel under three environments through single environment QTL analysis and joint analysis. The results of present study will be beneficial to elucidate the genetic basis of LA, fine map of QTL controlling maize LA, and design of a canopy ideotype at various canopy levels.

## Materials and Methods

### Plant materials and field experiment

The recombinant inbred line (RIL) population was developed by cross B73 and SICAU1212 as described previously (Yang *et al*., 2016; Yang *et al*., 2015a). The parent B73 with erect leaves is widely used as an elite line derived from the stiff stalk heterotic group and has been partly attributed to the changes in LA of maize varieties since 1970 (Russell *et al*., 1991), and another parent SICAU1212 with extremely expanded leaves derived from a waxy maize landrace Silunuo by continuously self-pollination 10 times and was cultivated at least 100 years ago (Tian *et al*., 2008). One hundred and ninety-nine RIL families were selected randomly from 325 RIL families which were developed in the previous study, and then used for QTL mapping in the present study.

The 199 RIL families along with the parent lines were phenotyped in three environments, which were located at Jinghong, Yunnan Province (21°57’N, 100°45E, elevation 551 m), in 2015 (15JH), Chengdu, Sichuan Province (30°43’N, 103°52’E, elevation 500 m), in 2016 (16CD), and Guiyang, Guizhou Province (26°29’N, 106°39’E, elevation 1277 m), in 2016 (16GY), respectively. The RIL families in each trial were planted with two replications by randomized complete block design. Fourteen plants of each family were cultivated in single-row plot with a planting density of 52,500 plants ha^−1^ in all environments. Row length was 3.0 m and row spacing was 0.67 m. Field management was the same as the standard cultivation management in accordance with growing season.

### Phenotype measurements and analysis

Five individuals from the middle of single plot were chosen to measure the leaf angle (LA) 10 days after flowering. Using the digital display protractor, we measured the LA of eight consecutive leaves below the tassel by manual refer to Hou *et al*.(2015). LA of the first leaf (the first leaf below tassel) was abbreviated as 1stLA, LA of the second leaf below the tassel was abbreviated as 2ndLA, etc. The phenotypic data of LA were determined as the average of each family from two replications in single environment. The variance components of genotype, environment, and G×E(genotype interacts with environment) were calculated by × SPSS Statistics version 20.0 software with general linear model (GLM) program (http://www.spss.com). Broad-sense heritability 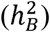 for each LA of eight leaves was estimated as 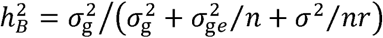, where 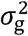 is the genetic variance, 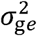 is the interaction variance between the genotype and environment, σ^2^ is the error variance, n is the number of the environments and r is the number of replications in each environment (Hallauer *et al*., 2010). Phenotypic correlation coefficients (r) between LA of eight leaves in each environment were also estimated by SPSS Statistics version 20.0 with Bivariate program.

### Linkage map and QTL mapping

The genetic linkage map used in this study derived from the linkage map of 253 RILs constructed in the previous study (Yang *et al*., 2016). In this study, the linkage map consisted of 260 molecular markers (106 SSR and 154 InDel markers). The number of markers located on each chromosome ranged from 18 to 35, covering about 88.95% of the maize genome. The total length of the linkage map was about 1133.57 cM and the average interval length between adjacent markers was 4.36 cM. The map was drawn by MapChart software version 2.3 (Voorrips, 2002). QTL for each LA of eight leaves were detected by including composite interval mapping (ICIM) model (Li *et al*., 2008; Li *et al*., 2007) using the QTL IciMapping software version 4.0 (http://www.isbreeding.net/) in single environment. The parameters were that the walking speed was 1.0 cM, the probability in the stepwise regression was set to 0.001,and threshold LOD scores were determined by 1,000 permutations and a type I error was set at P = 0.05. The joint mapping, epistatic interaction and QTL by environment interaction (QEI) detection were identified by the mixed-model-based composite interval mapping (MCIM) (Wang *et al*., 1999) using the QTLNetwork software version 2.1 (Yang *et al*., 2008). The testing window size, walk speed and filtration window size of the genome scan configuration were set to 10, 1 and 10 cM, respectively, and significant QTL were also determined by 1,000 permutations as *P* = 0.05. The name of QTL, such as *qLA1.1*, was assigned as ‘q’ followed by ‘LA’, ‘maize chromosome on which the corresponding QTL is located’, ‘.’, and ‘serial number of QTL’. These QTL for LA were deemed to be a same QTL when the confidence intervals of such QTL were overlapped or shared one marker. Additionally, the QTL with PVE > 10% was declared as the major QTL. The QTL was considered as stable QTL that was identified in two or three environments.

## Results

### Phenotypic variation in LA of eight consecutive leaves

The phenotypic values of LA were analyzed in the RIL families and their parent lines cultivated in three distinct habitats (Table 1). All eight leaves tested in the parent B73 displayed a relatively vertical angle (less than 45°), and the higher the leaf position was, the smaller the LA was; whereas parent SICAU1212 had more horizontal leaf orientations (more than 45°); the difference among LA of eight leaves between these two parents could be also observed in Fig.1 in the previous study (Yang *et al*., 2016). It was obvious that each LA of eight leaves in B73 was significantly different from that in SICAU1212 (*P* < 0.01). In addition, all LA of eight leaves showed a normal distribution with transgressive segregation in three environments, suggesting quantitative genetic control (Supplementary Figure S1). The ANOVA analysis indicated that genotype, environment, and G × E interactions within the RIL population were highly significant (*P* < 0.001) different in all LA of eight leaves (Table 2); Moreover, replications of all LA of eight leaves except for the 8thLA were non-significant (*P* < 0.05); hence, the average of the two replications of each RIL family in one environment was used to QTL mapping. Broad-sense heritability 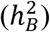 for these LA of eight leaves were relatively high, which ranged from 79.47% to 83.46% (Table 2), indicating that much of the LA variation in the RIL population was genetically controlled.

**Figure 1.**
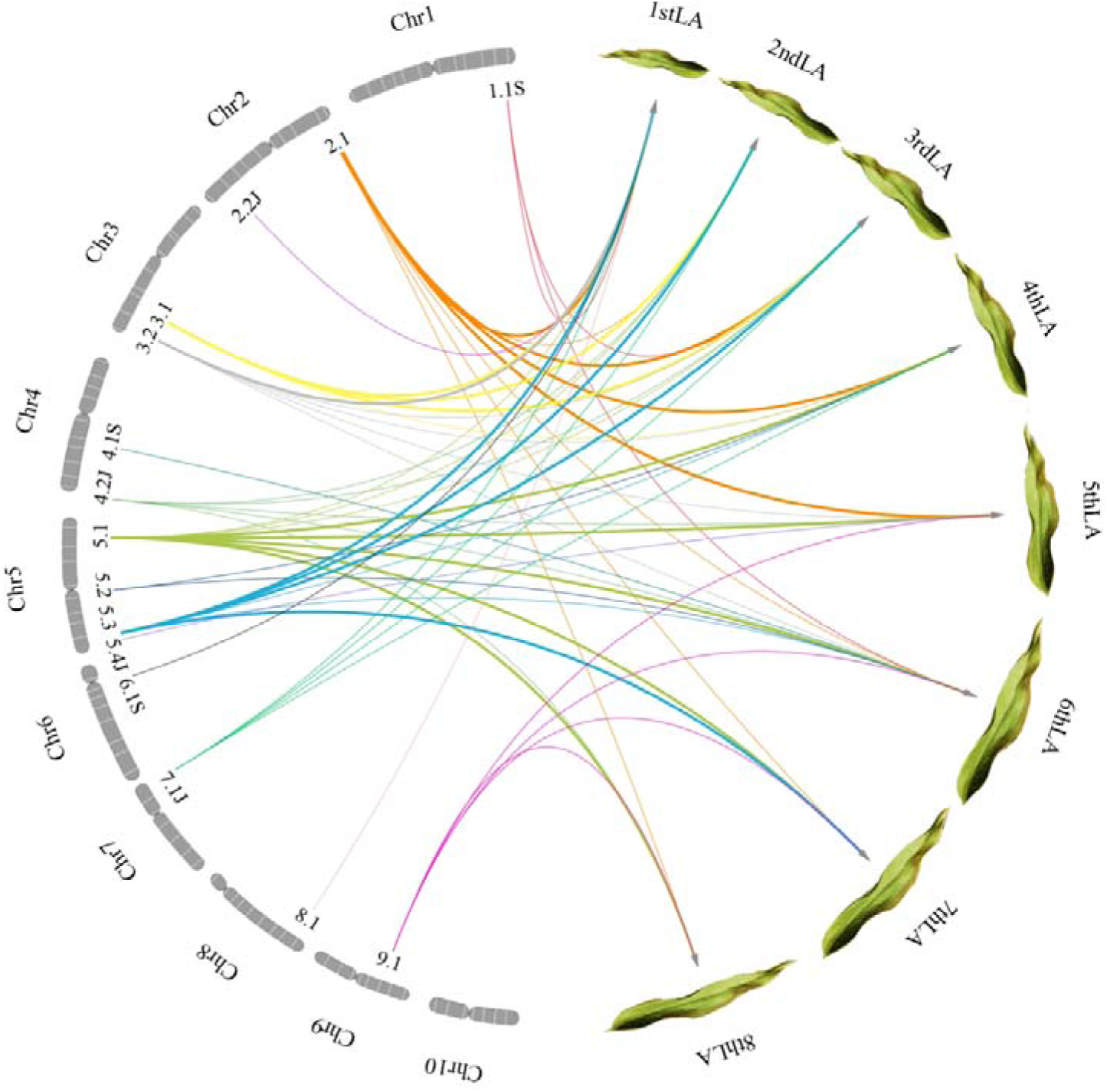
Network diagram of identified QTL controlling eight LAs. The 1.1 under chromosome 1 (Chr1) represents QTL *qLA1.1*, and so forth. The capital letters S and J represent the QTL detected only in single-environment and joint analysis, respectively. The arrowhead lines mean that the QTL control the corresponding LA. Different colored lines represent different QTL. The thick lines represent QTL with PVE > 10%; fine lines represent QTL with PVE ≤ 10%. When QTL were identified in more than one environment or in both single-environment and joint analysis, the QTL with the highest value of PVE was selected.

**Table 1.**
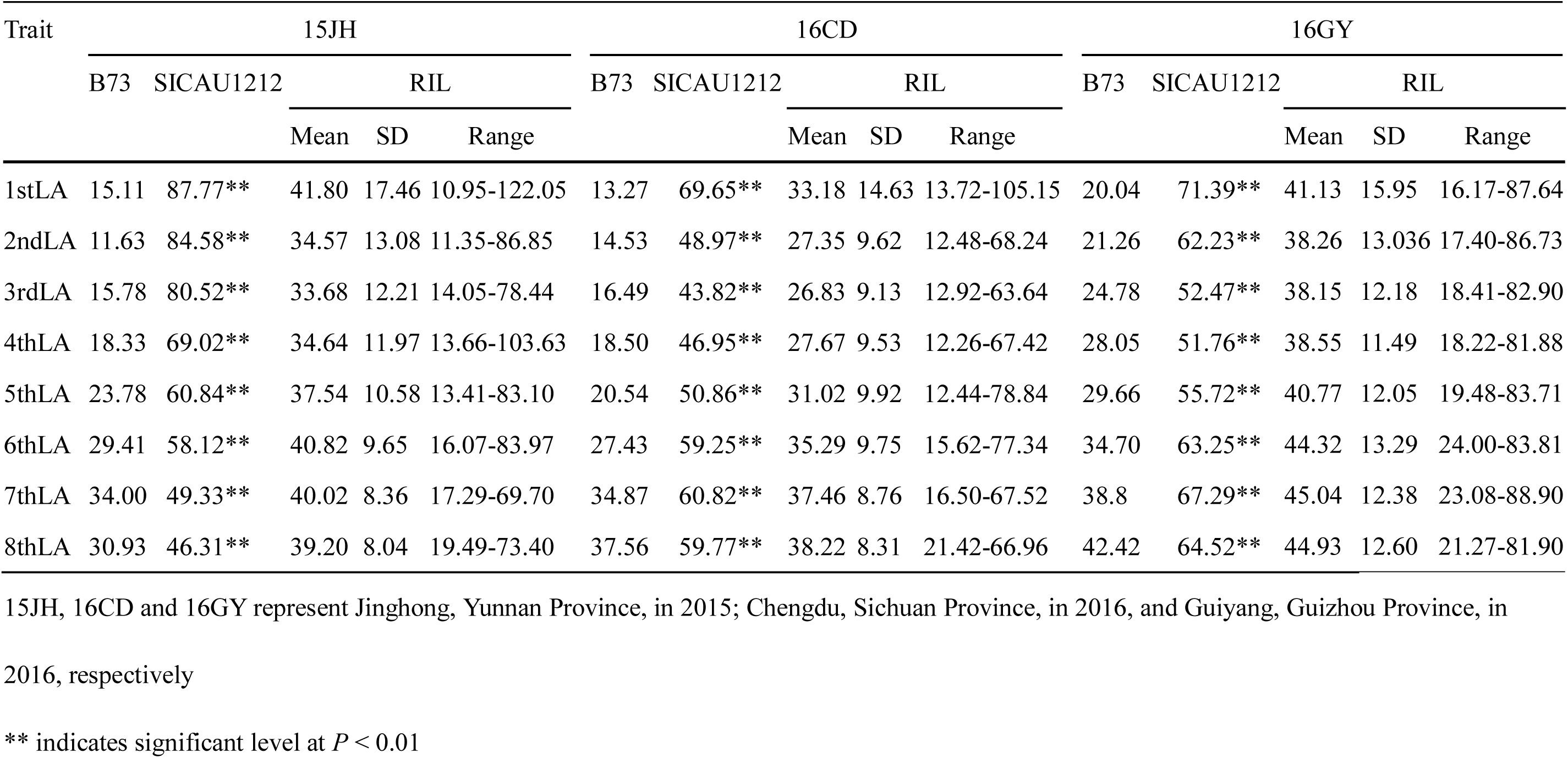
Phenotypic performance for leaf angle of the RIL population and parents in three environments

**Table 2.** Analysis of variance (ANOVA) of LA in the RIL population in three environments

| Source of variation | Mean square |  |  |  |  |  |  |  |
| --- | --- | --- | --- | --- | --- | --- | --- | --- |
|  | 1stLA | 2ndLA | 3rdLA | 4thLA | 5thLA | 6thLA | 7thLA | 8thLA |
| Environment (E) | 9966.08*** | 12394.80*** | 13288.85*** | 12193.37*** | 9536.30*** | 7790.14*** | 6213.45*** | 6187.40*** |
| Genotype (G) | 705.19*** | 357.06*** | 292.89*** | 291.92*** | 267.25*** | 269.66*** | 238.37*** | 240.52*** |
| G × E | 349.42*** | 197.47*** | 180.40*** | 198.24*** | 168.31*** | 181.75*** | 170.71*** | 163.32*** |
| Replication | 2.76 | 23.50 | 180.95 | 79.70 | 29.56 | 129.28 | 3.47 | 180.63* |
| Error | 139.70 | 64.72 | 62.40 | 55.98 | 52.13 | 50.75 | 46.40 | 39.71 |
| $h_B^2$ | 83.46 | 82.33 | 80.59 | 79.47 | 80.49 | 79.62 | 78.67 | 79.75 |

$h_B^2$ , the broad-sense heritability
\*, \*\* and \*\*\* indicate significant level at $P < 0.05$ , $P < 0.01$ and $P < 0.001$ , respectively

### Correlation analysis

The phenotypic coefficients between LA of eight leaves from three environments show highly significance in RIL families, and the correlation coefficients varied from 0.449 to 0.907 (Supplementary Table S1). Overall, the values of correlation coefficients between corresponding LA of two leaves in the three environments were roughly equal, and all had significant positive correlations. However, for the two leaves, the farther apart the two leaves, the smaller the correlation coefficient of LA; thus, the correlation coefficients were the highest between adjacent two leaves.

### Single environment QTL analysis and joint analysis

Using inclusive composite interval mapping, a total of 56 putative QTL for LA of eight leaves were identified in three environments, distributed on 10 chromosomes except chromosome 7, with each QTL accounting for 5.62% - 20.14% of the phenotypic variation (Table 3, Supplementary Figure S2). Among them, 52 QTL could be divided into 7 common QTL since they overlapped or shared a common marker. Of these 7 common QTL, four QTL (*qLA2.1*, *qLA3.1*, *qLA5.1*, *qLA9.1*) were identified in two or three environments, the remaining QTL were only detected in one environment. For a LA of specific leaf (1stLA to 8thLA), 1 to 5 QTL were identified for LA in single environment, together explained 8.52% - 48.11% of the phenotypic variation. In addition, four QTL including *qLA4.1*, *qLA5.2*, *qLA6.1* and *qLA8.1* that affected the variation of LA of one leaf; while the others 7 QTL including *qLA1.1*, *qLA2.1*, *qLA3.1*, *qLA3.2*, *qLA5.1*, *qLA5.3* and *qLA9.1* that contributed to the variation of LA of 3-8 leaves. It is worth noting that *qLA5.1* affected LA of all eight leaves, followed by *qLA2.1* controlling LA of seven leaves; although both *qLA3.1* and *qLA9.1* affected LA of four leaves, *qLA3.1* affecting LA of upper four consecutive leaves (1stLA to 4thLA) and *qLA9.1* affecting LA of lower four consecutive leaves (5thLA to 8thLA); furthermore, *qLA1.1*, *qLA3.1* and *qLA5.3* affected LA of 3-5 non-consecutive leaves. All QTL except for *qLA6.1* had a negative additive effect in the single-environment analysis, suggested that the alleles derived from SICAU1212 for detected QTL increased the value of LA.

**Table 3.**
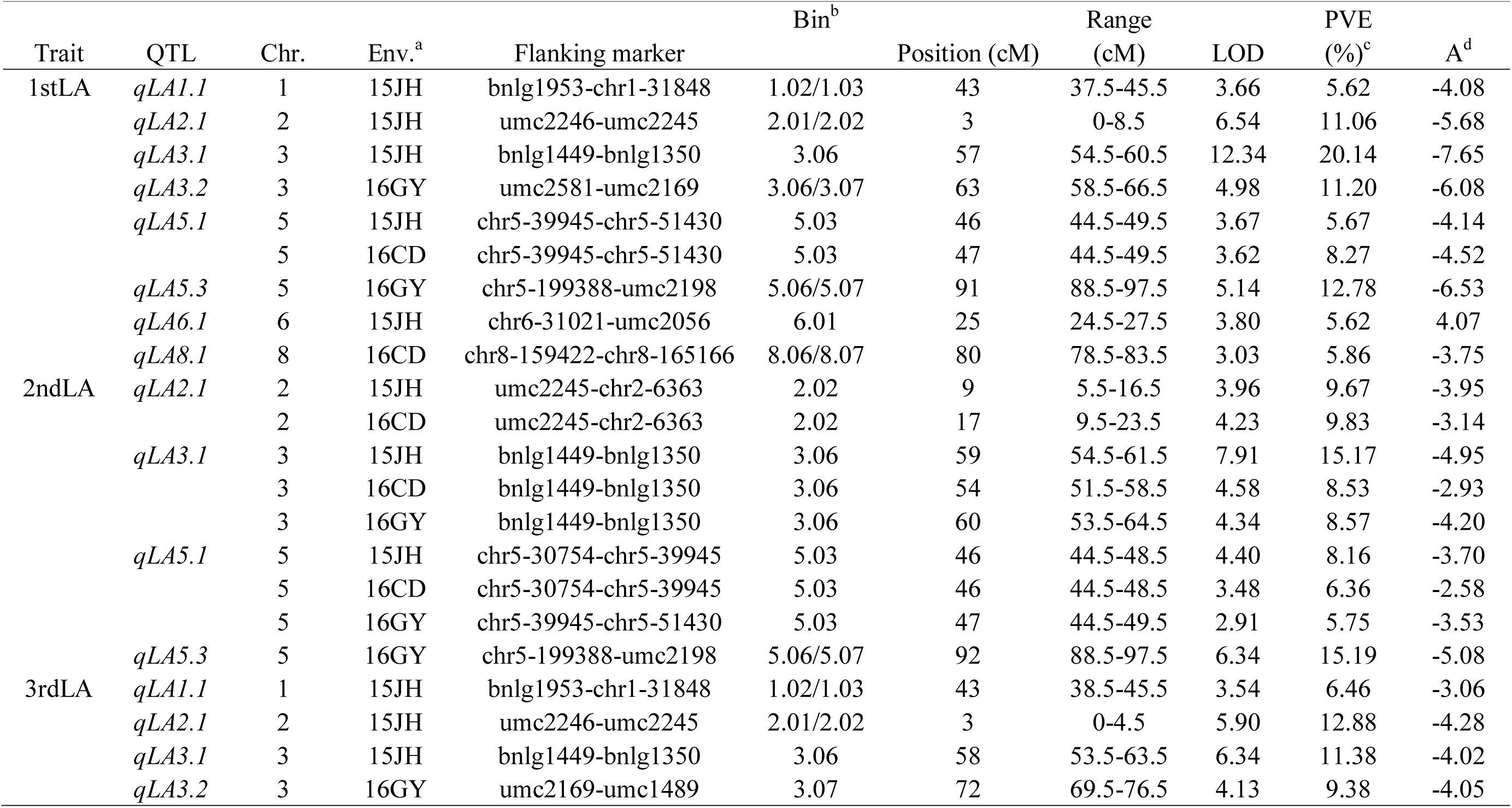
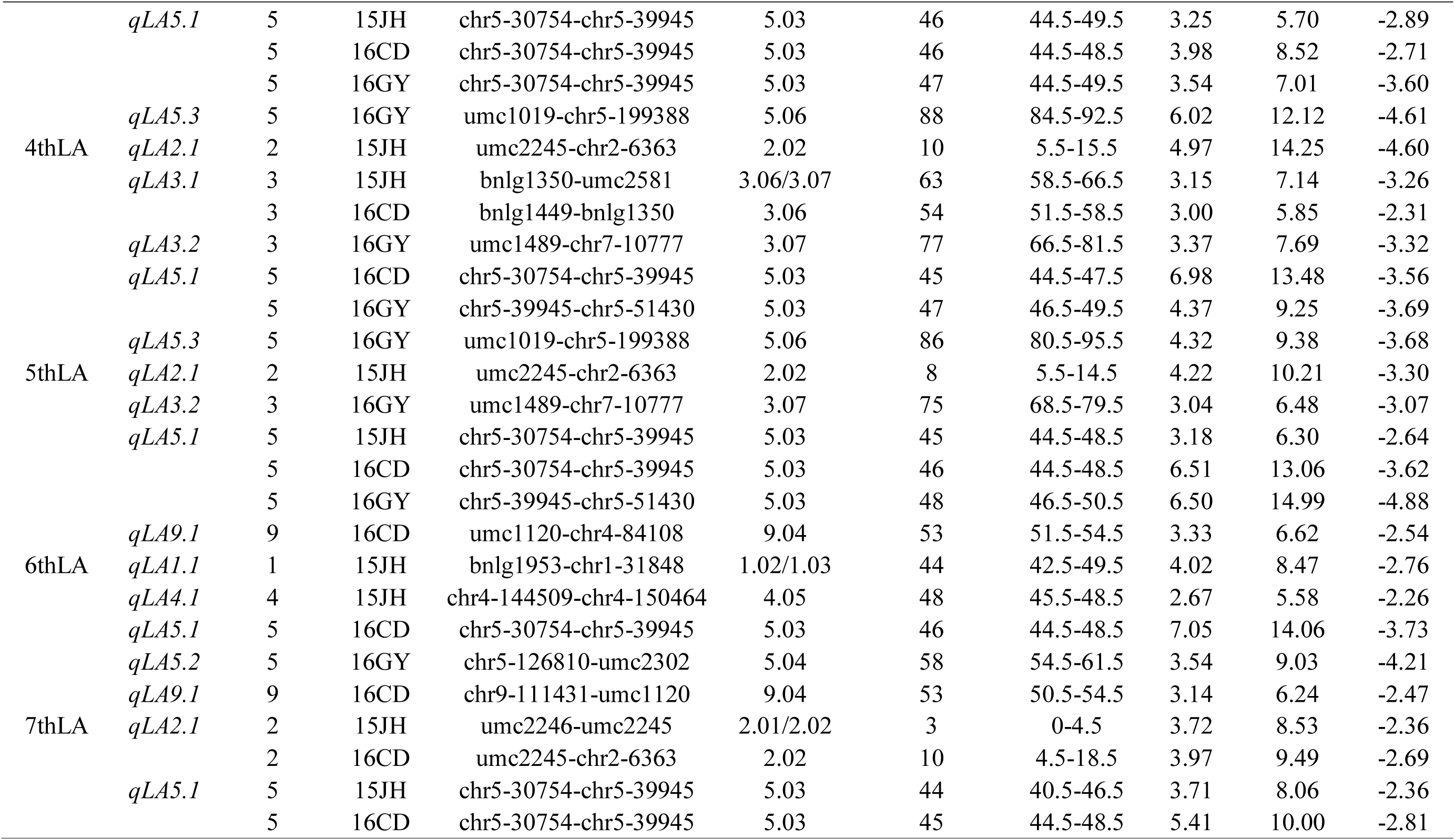
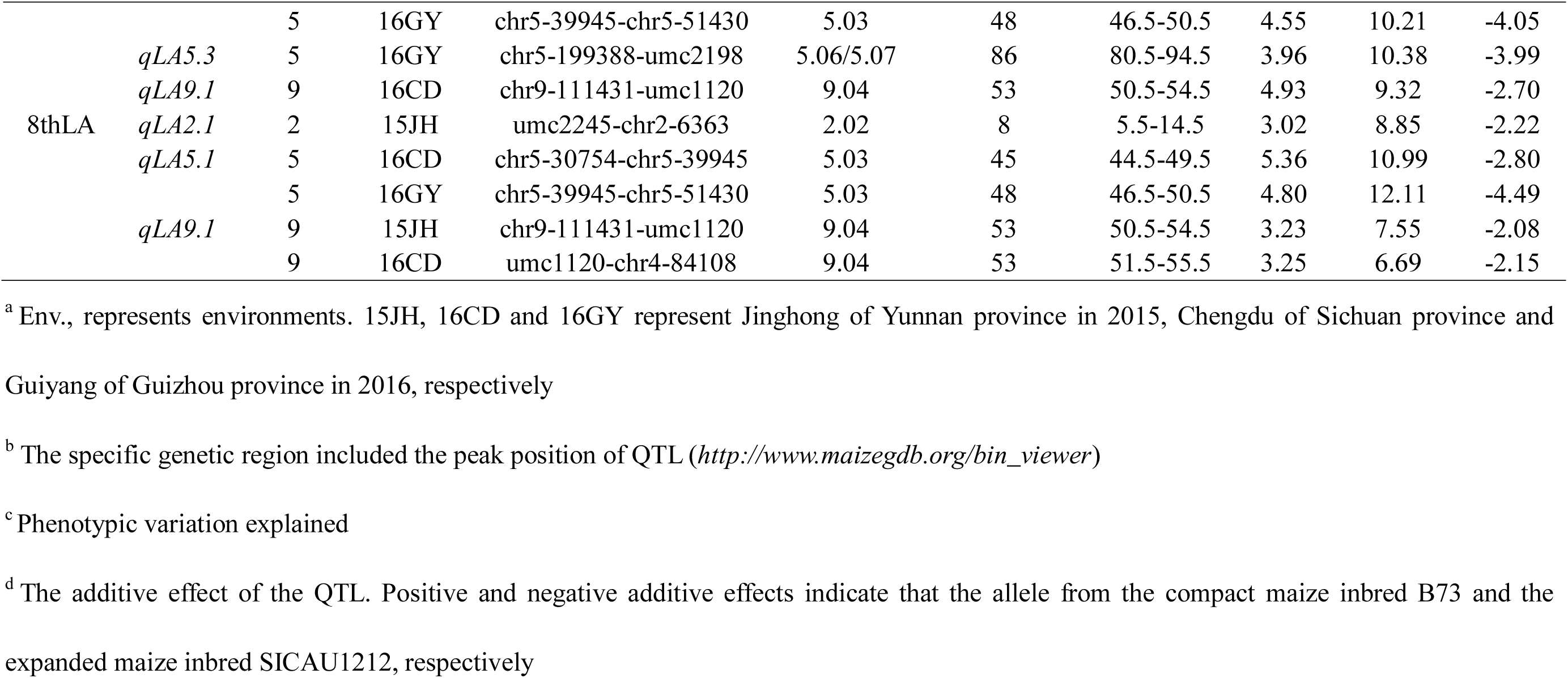
Putative QTL for LA in the RIL population through single-environment QTL mapping

Meanwhile, forty-four significant QTL for LA of eight leaves were detected in joint analysis, distributed on 10 chromosomes except for chromosome 1 and 10, with single QTL explaining 1.29% - 10.79% of the phenotypic variation (Table 4, Supplementary Fig. S2). Of these QTL, 41 QTL could be classified as 9 common QTL, each common QTL affecting LA of 2-8 leaves; 7 common QTL were identical to QTL identified in single-environment analysis; while 4 QTL were only identified in joint mapping, including *qLA2.2*, *qLA4.2*, *qLA5.4* and *qLA7.1.* All QTL except for *qLA2.2* and *qLA4.2* had a negative additive effect, suggested that most of the alleles with a contribution on increasing LA were segregated from SICAU1212 with expanded plant architecture.

**Table 4.**
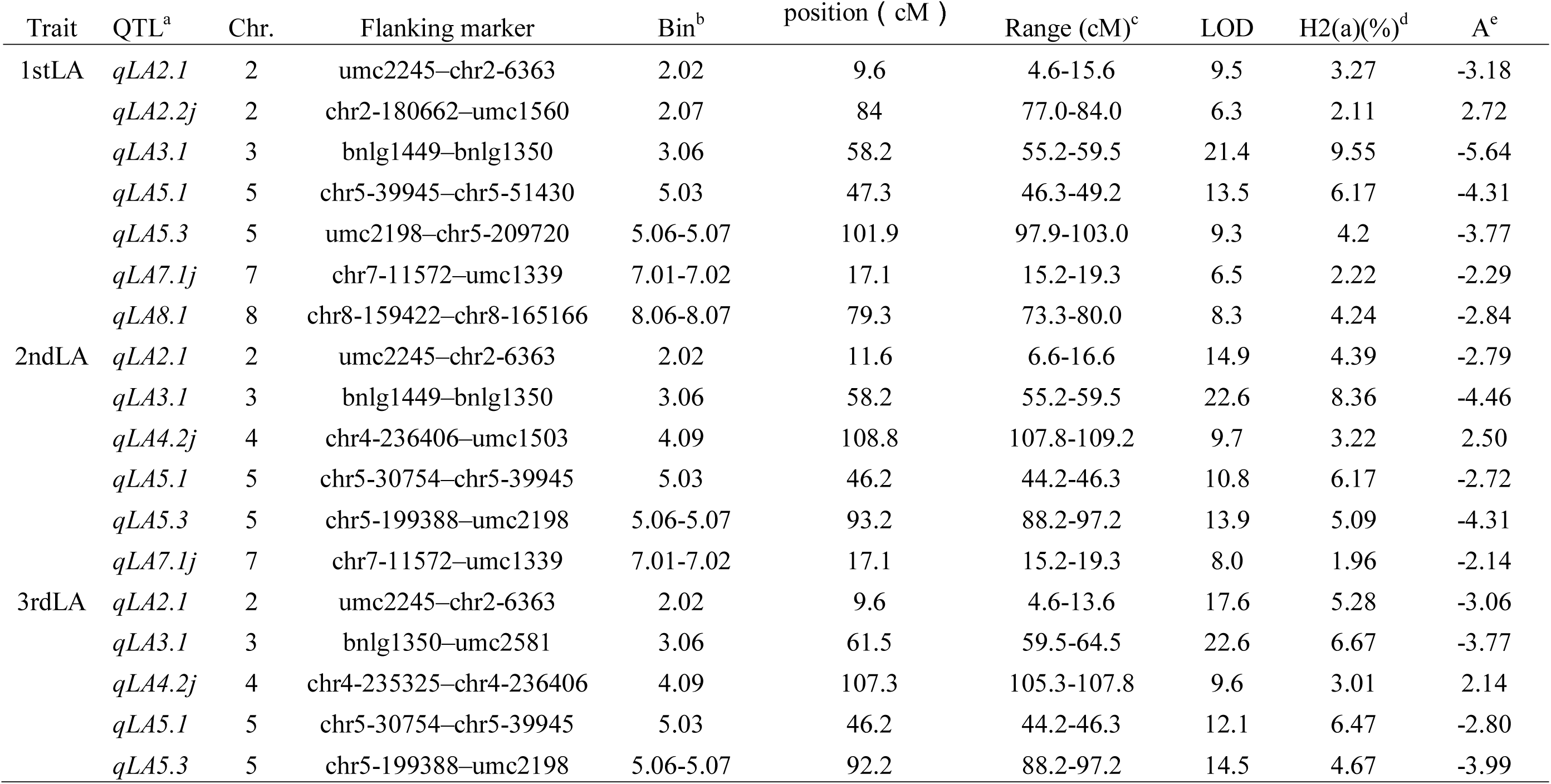
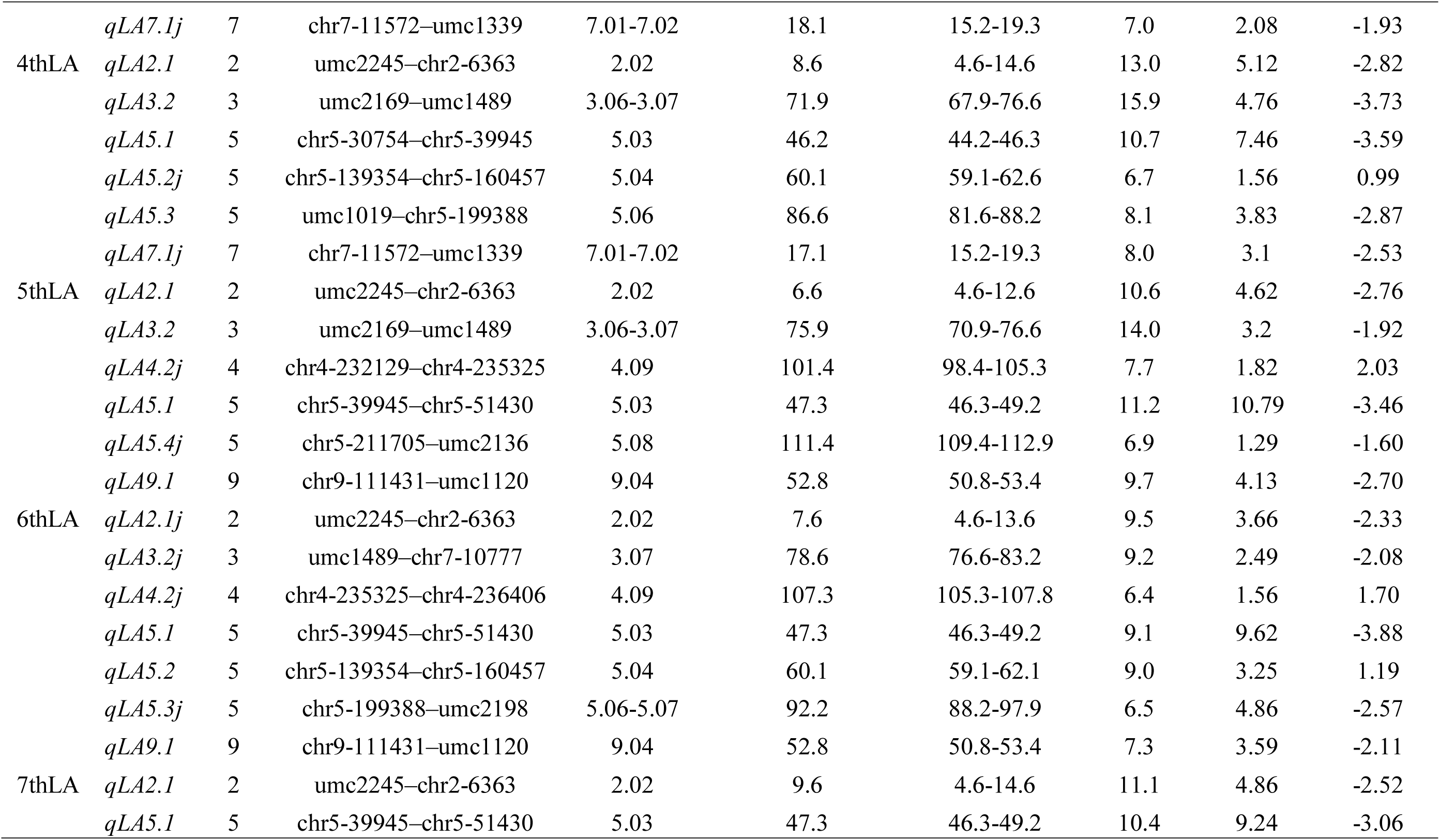
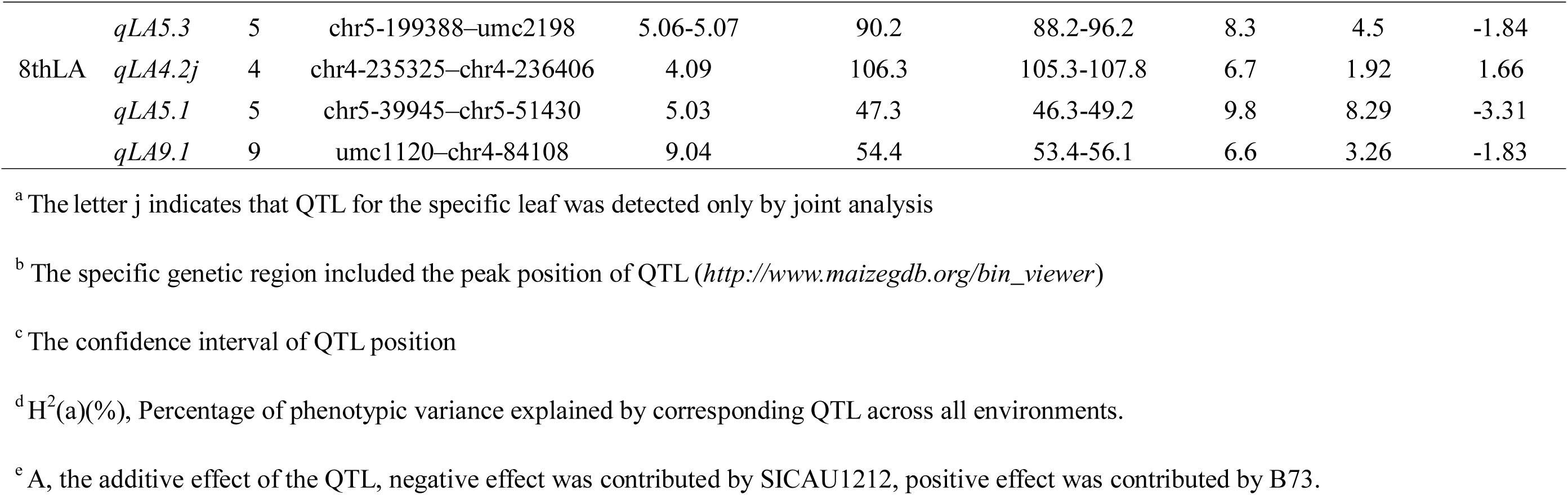
QTL for eight LAs detected in joint analysis across three environments

### QTL × Environment (QE) interactions

Four QTL were involved in significant QTL × environment interaction (QEI) (Table 5) through joint analysis. Three QTL of them were common QTL (*qLA5.3*) that affected the LA of the 2nd, 3rd and 7th leaves simultaneously; the additive × environment interactions for LA were responsible for 1.89% - 2.46% of phenotypic variation. The other QTL (*qLA5.2*) was associated with the LA of the 6th leaf, and the effect of additive × environment interaction was 1.50%.

**Table 5.** QTL × Environment (QE) interactions for LAs identified in the RIL population using QTLNetwork

| Traits | QTL | Marker interval | AE1(15JH) | AE2(16CD) | AE3(16GY) | H<sup>2</sup> (ae)(%) |
| --- | --- | --- | --- | --- | --- | --- |
| 2ndLA | *qLA5.3* | chr5-199388–umc2198 |  | 1.90<sup>*</sup> | -2.79<sup>**</sup> | 2.05 |
| 3rdLA | *qLA5.3* | chr5-199388–umc2198 |  | 1.92<sup>**</sup> | -2.32<sup>**</sup> | 1.89 |
| 6thLA | *qLA5.2* | chr5-139354–chr5-160457 |  |  | -1.29<sup>*</sup> | 1.50 |
| 7thLA | *qLA5.3* | chr5-199388–umc2198 |  |  | -2.07<sup>**</sup> | 2.46 |

AE is the additive by designated environment interaction effect
H<sup>2</sup> (ae)(%) is contribution rate of additive by environment interaction
\*, \*\* and \*\*\* indicate significant level at $P < 0.05$ , $P < 0.01$ and $P < 0.001$ , respectively

### Epistatic interaction

A total of twenty epistatic interactions with additive-by-additive effects were identified for LA of eight leaves with individual interaction accounting for 0.39% - 3.54% of the phenotypic variation (Table 6). These interactions could be divided into three types of epistatic interactions, including interactions between the genetic regions of identified QTL, between significant QTL and non-significant QTL region, and between non-significant QTL regions. However, the number of leaves affected by each epistatic interaction was different, ranging from 1 to 4. For instance, the epistatic interaction between *qLA5.3* and *qLA7.1* affected four leaves (1stLA to 4thLA), and the epistatic interaction between the marker intervals of chr9-90756–mmc0051 and chr10-77445–umc1336 only affected the 2ndLA. Additionally, the number of epistatic interactions of various leaves (1stLA to 8thLA) was different, varying from 0 to 6. For example, five epistatic interactions were identified for the 2ndLA, while no epistatic interaction was identified for the 8thLA.

**Table 6.**
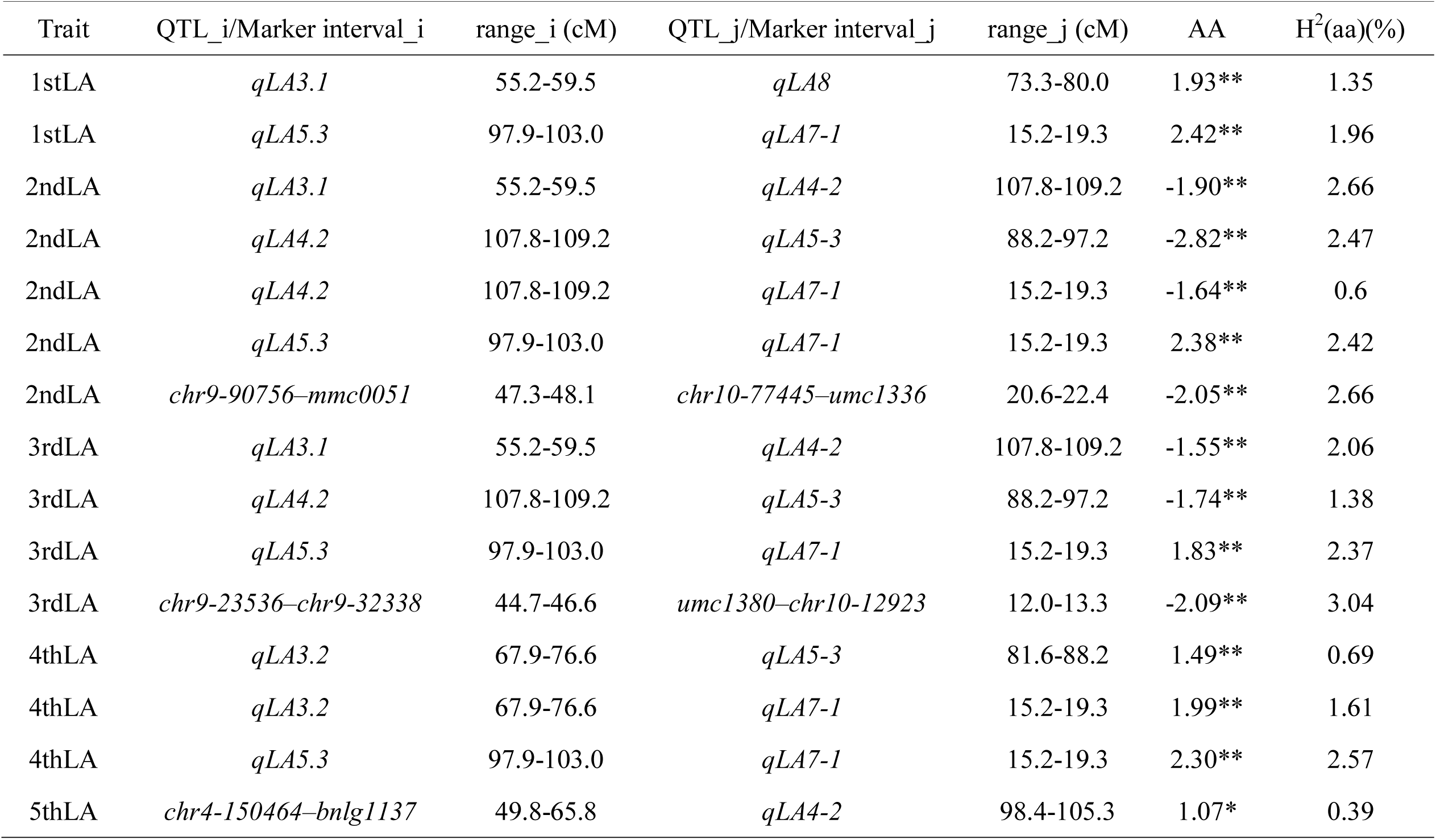
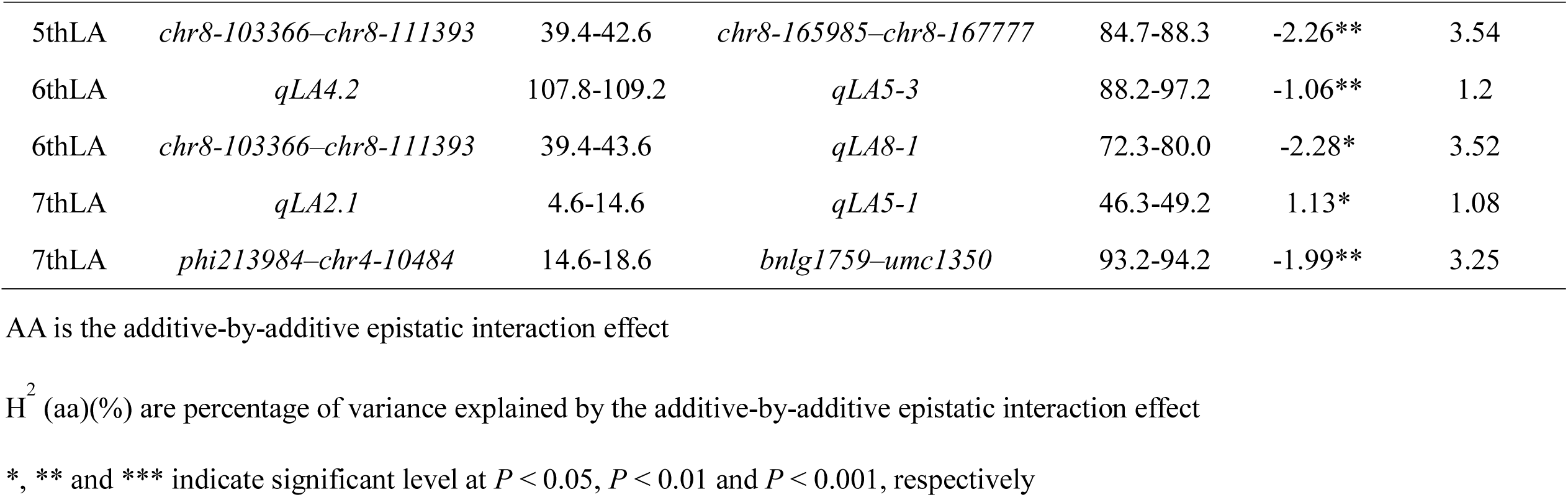
Digenetic epistatic QTL for LA identified in the RIL population across three environments

## Discussion

### Comparison of the mapped QTL with previously identified QTL and genes

In this study, we performed QTL mapping for LA of eight consecutive leaves below the tassel, a total of 56 putative QTL were mapped in single environment analysis and 44 QTL were identified in joint analysis; Among them, there were 9 common QTL because they could affect more than one LA or were identified in more than one environments, which were hotspot regions for LA distributed on chromosome 1, 2, 4, 5, 7, 8 and 9 (Supplementary Table S2; Supplementary Figure S2). Meanwhile, comparing the QTL in this study with QTL previously identified, we found that all QTL were consistent with QTL identified in at least one previous study (Supplementary Table S2). For instance, QTL *qLA1.1* on chromosome 1 (∼23.92-31.84 Mb, bin 1.02/1.03) was consistent with that of 12 previous studies, harboring the gene *drooping leaf1* (*drl1*) (∼26.77 Mb). *Drl1* encodes a transcription factor belonging to YABBY family, which is required for leaf development and for proper leaf patterning (Strable *et al*., 2017). In addition, *grassy tillers 1* (*gt1*) (∼23.62 Mb) is close to *qLA1.1*, which belongs to class I homeodomain leucine zipper gene family, and regulates the tillers of maize and responds to shade signal (Whipple et al. 2011;). QTL *qLA2.1* on chromosome 2 (∼3.08-6.36 Mb, bin 2.01/2.02) was agree with that of 12 previous studies, harboring the gene *liguleless1* (*lg1*) (∼4.23 Mb); *lg1* mutant lacks a ligule or an auricle, resulting in produce much more erect leaves (Becraft et al. 1990; Becraft and Freeling 1991; Sylvester et al. 1990); *lg1* encodes a SPL transcription factor, along with its homologs *TaSPL8* in wheat and *OsLG1* in rice, regulating auxin and brassinosteroid signaling, thus leaf angle (Ishii et al. 2013; Kong et al. 2017; Lewis et al. 2014; Liu et al. 2019; Tian et al., 2019). QTL *qLA3.1* on chromosome 3 (∼135.52-182.12 Mb, bin 3.06) overlapped with the QTL detected by Lu et al. (2007) and Dzievit et al. (2019), which contained the gene *liguleless2* (*lg2*) (∼179.38) encoding a bZIP transcription factor that function in the narrowed region of ligule and auricle (Walsh et al. 2010). QTL *qLA5.1* on chromosome 5 (∼30.75-51.43 Mb, bin 5.03) was consistent with that of 7 previous studies; however, no gene has been cloned for LA variation in this QTL region; since *qLA5.1* could affect LA of all eight leaves, and explained 5.67-14.99% of the phenotypic variation across three environments, it is worth fine mapping for it. Additionally, several QTL, such as *qLA2.2*, *qLA3.2*, *qLA4.1* and *qLA5.4*, were overlapped with the QTL mapped in 2 out of 29 previous studies listed in Supplementary Table S2, which may be rare loci for LA variation. However, we did not identify LA QTL on chromosome 2 (bin 2.05) containing *UPA2* (*ZmRAVL1*), the reason probably is that *UPA2* is rare allele from teosinte and was lost during maize domestication as mentioned in the report (Tian et al. 2019).

### The phenotype and genetic architecture of the eight leaf angles exhibit extensive diversity

Leaf angle is a crucial factor affecting the plant architecture and associated with yield indirectly (Liu et al. 2019). The range of variation in LA at different leaves in individual maize plant was relatively large. As we observed, the LA of the leaves at various nodes was different, for instance, the LA changed 1.6-fold and 2.6-fold among eight leaves in parental lines SICAU1212 and B73 in this study, respectively. In addition, the average LA of three leaves above the ear, three ear leaves and three leaves below the ear were 22.9°, 32.8° and 41.9° in the high-yielding maize hybrid Pioneer 335, respectively; thus, there was about 2.0 fold change between the LA of uppermost and lower leaves (Ma *et al*., 2014b). These rich variations in LA of a single plant provide a possibility that we can modulate the LA of leaves in various nodes. Moreover, the range of variation of LA in maize inbred lines is also very large, which is the second highest degree of variation traits and is second only to the tassel branch number (Pan *et al*., 2017). In this study, the LA at eight leaves in SICAU1212 were more than 45°, which were about 1.5 - 5.8 fold larger than that in B73. The LA of upper leaves of 26 maize inbred lines that used to construct NAM population variated from approximately 30° to 80° (Tian *et al*., 2011). Similarly, the middle leaf angle (MLA) ranged from 23.8° to 38.7° in 14 elite maize inbred lines that used to develop the ROAM population (Pan *et al*., 2017). Using these maize inbred lines with a large variation of LA, we may breed various maize hybrids to adapt different ecological environments and plant density. In view of the extensive and continuous phenotypic variation of LA, there may be a large number of loci controlling LA and each locus just explains a small part of variation (Pan *et al*., 2017).

On the other hand, QTL mapping for LA of eight leaves indicated that there were different sets of QTL that controlled LA at various leaves, revealing a distinct genetic architecture of the LA, as follow (i) although the major QTL with PVE > 10% were detected for all eight LA across three environments, the number of major QTL for each LA was difference, ranging from 1 (6thLA and 8thLA) to 4 (1stLA); (ii) the number and effect of epistatic interaction for each LA were difference; No epistatic interaction was detected for the 8thLA, while 2 to 5 epistatic interactions were identified for the other seven LA, explaining 3.31% - 10.81% of the phenotypic variation; (iii) LA of four leaves (2ndLA, 3rdLA, 6thLA and 7thLA) were affected by QTL × environment interactions (QEI), whereas no QEI was identified for the remaining LA of four leaves. From these comparative analyses, we can infer that the 8thLA was mainly affected by major QTL plus minor QTL; the 1stLA, 4thLA and 5thLA were mainly controlled by major QTL, minor QTL and epistatic interactions; and the remaining four LA (2ndLA, 3thLA, 6thLA and 7thLA) were affected not only by major QTL, minor QTL and epistatic interactions, but also the environment. But, it is important to note that, for LA of eight leaves, the broad-sense heritability ranged from 78.67% to 83.46%, which was relatively high and almost equal. However, QTL for each LA, which was identified in single environment, just explained 8.52% - 48.11% of the phenotypic variation, revealed that parts of phenotype variation could not be explained. The reason may be that numbers of QTL with minor effect and interactions were not identified.

### Possibility of manipulating LA at canopy-wide levels

The smart canopy concept was proposed that improvement of light harvesting and metabolic features of the leaves interacting cooperatively at the canopy level to maximize the potential yield (Ort *et al*., 2015; Yang *et al*., 2016). To engineer the plant in a “smart” canopy, the LAs of plant should be changed in the way that leaves of upper canopy are vertical (small leaf angle) and leaves of lower canopy are horizontal, thus permitting more light reach the canopy(Yuan et al. 2001; Ort *et al*., 2015). However, the genetic basis of leaf angle at upper and lower canopy levels was not well known. Although many studies were conducted to QTL mapping for LA, a fewer leaves (1-4 leaves) were selected to phenotyping the LA, including the second leaf below tassel (Feng et al. 2015), the third leaf below tassel (Yu et al. 2006), four leaves above the uppermost ear (Ding et al. 2015), the ear leaf (Ku et al. 2012; Ku et al. 2010; Shi et al. 2017), the first leaf below the primary ear (Hou et al. 2015), the second leaf below the ear (Dzievit et al. 2019), and so on. Hence, it is hard to differentiate whether a QTL controls LA of one leaf or multiple leaves (Dzievit et al. 2019; Mantilla-Perez and Salas Fernandez 2017). In this study, QTL mapping was conducted for LA of eight consecutive leaves below tassel, and the QTL identified could answer the question partly mentioned above, as follow, QTL like *qLA5.1* could control LA of all eight leaves, which provides the possibility that regulation of leaves at the whole canopy; QTL such as *qLA3.1* could affect LA of the upper four leaves, leading to the potential to modulate LA in the upper canopy; while QTL *qLA9.1* could affect LA of the lower four leaves, leading to the potential to regulate LA in the lower canopy; moreover, QTL including *qLA4.1*, *qLA6.1*, *1stLA* only affect LA of one specific leaf, which is help to regulate LA of the leaf we need, not the others. Together, these various QTL not only provides the possibility that regulation of the LA at different canopy levels, but also help to unlock a door for further dissection of LA in maize.

## Abbreviation

LA: Leaf angle
1stLA: First leaf angle
2ndLA: Second leaf angle
3thLA: Third leaf angle
4thLA: Fourth leaf angle
5thLA: Fifth leaf angle
6thLA: Sixth leaf angle
7thLA: Seventh leaf angle
8thLA: Eighth leaf angle
QTL: Quantitative Trait Locus

## Acknowledgements

The authors are grateful to the National Basic Research Program of China (the “973” project, Grant No. 2014CB138203), the State Key Laboratory of Grassland Agro-Ecosystems, China (SKLGAE201509) and the National Natural Science Foundation of China (31101161).

## Supplementary data

**Table S1.** Phenotypic correlation coefficients between eight LAs across three environments

**Table S2.** Summary of QTL identified in this study and previous studies

**Figure S1.** Frequency distribution of LA from eight nodes under three environments

**Figure S2.** Genetic linkage map of the RIL population and locations of QTL for eight LAs in single-environment QTL mapping and joint analysis

**Additional file 1** Genotypes of the RIL population

**Additional file 2** Phenotypes of leaf angle of eight consecutive leaves in the RIL population across three environments

## Compliance with ethical standards Conflict of interest

The authors declare that they have no conflict of interest.

