## Supplementary Figure S1-S2 for "Identification of QTL for leaf angle at canopy-wide levels in maize"

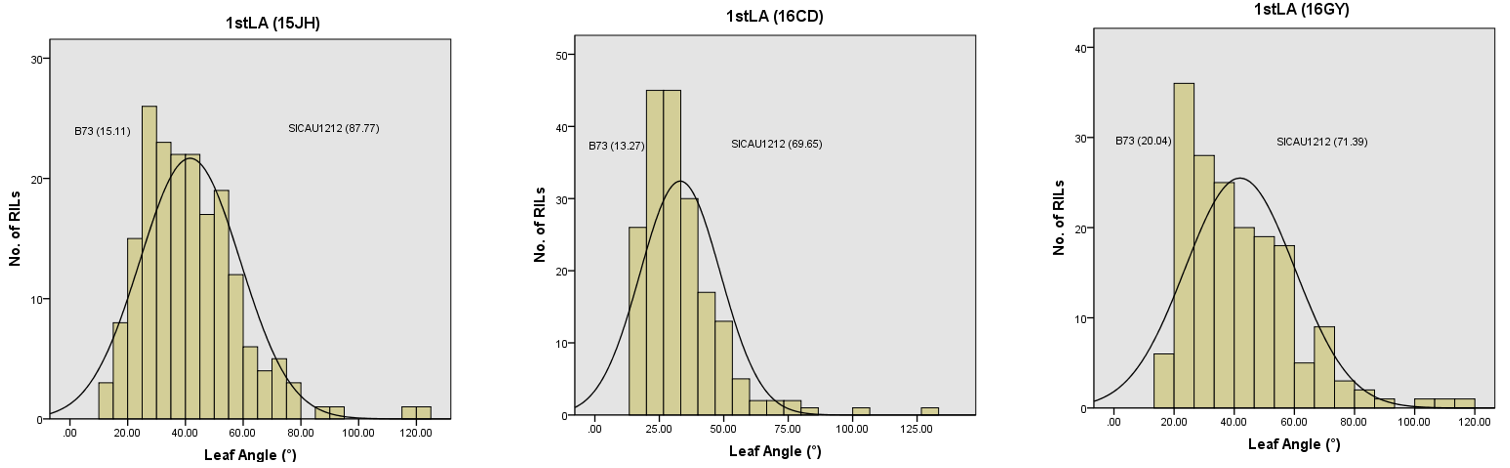

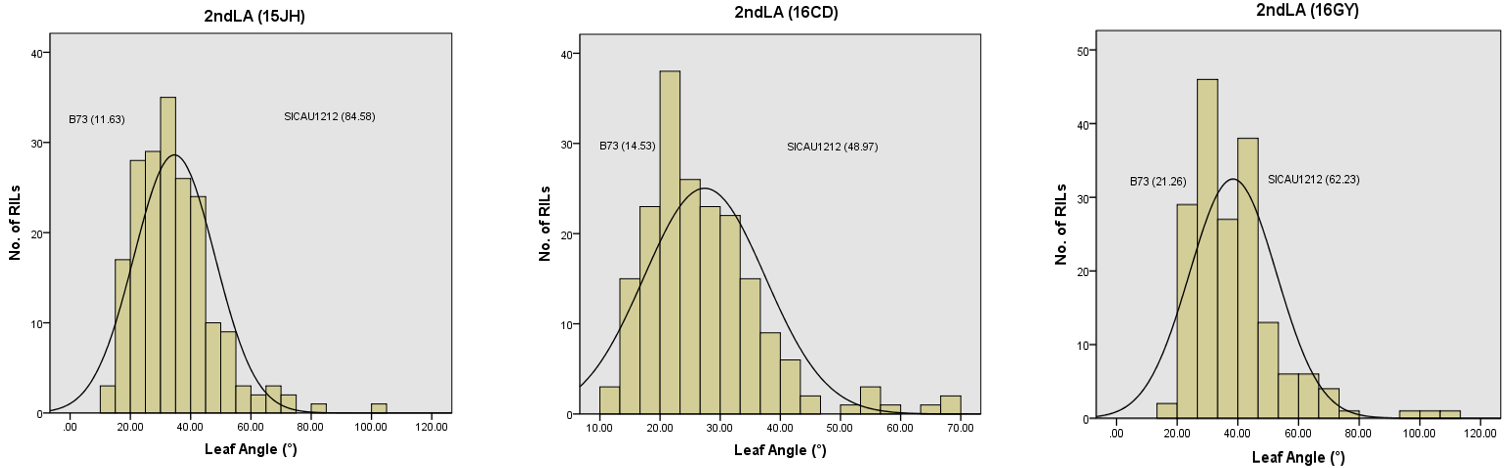

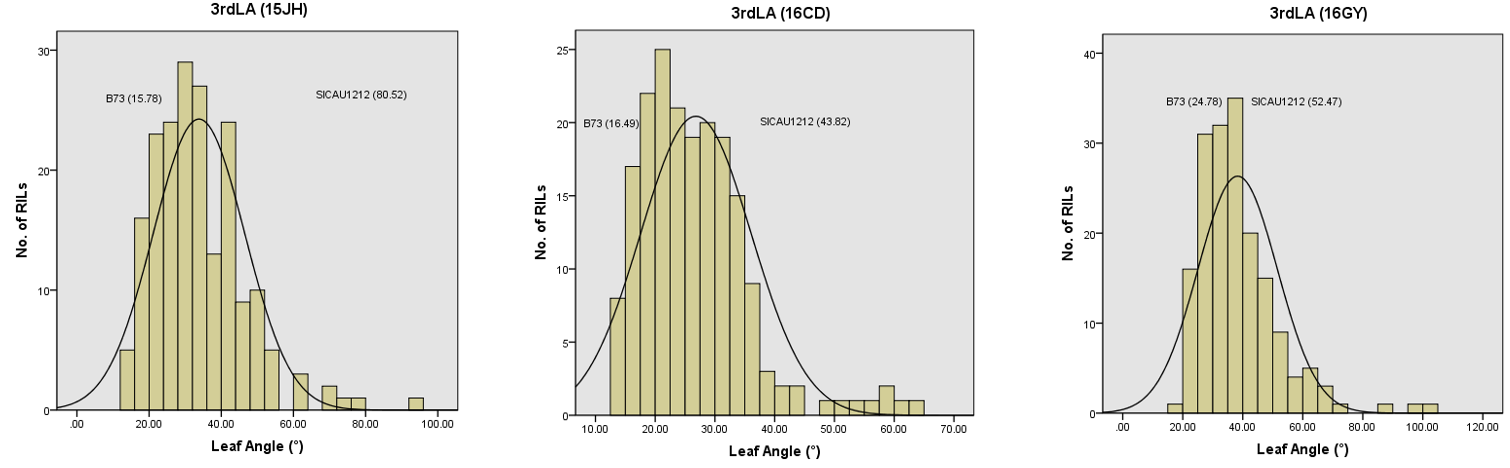

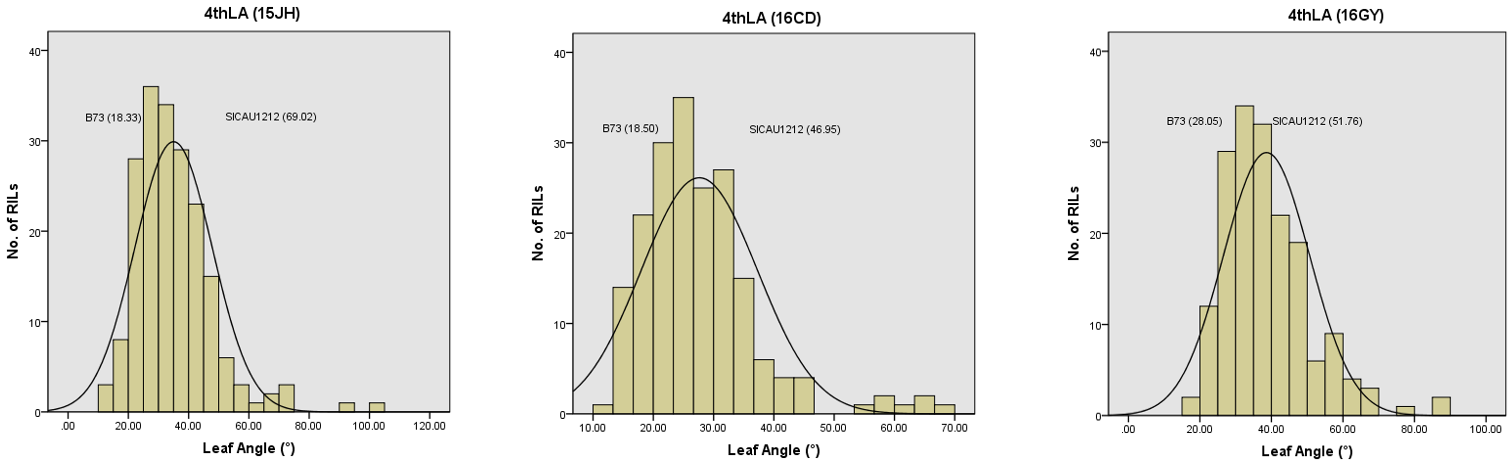

**Figure S1 continued**

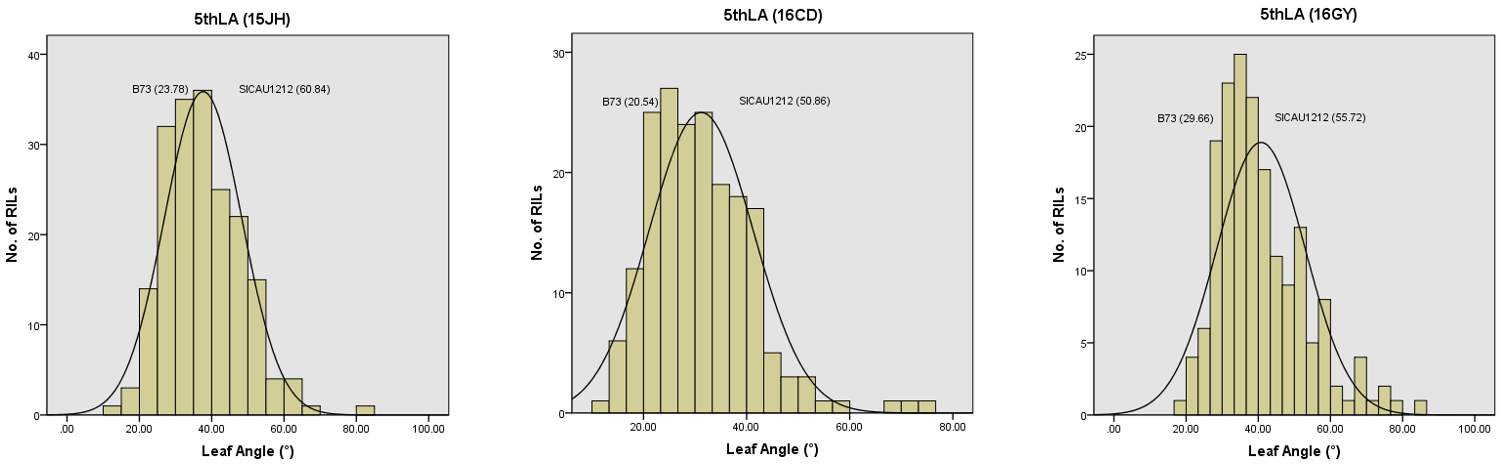

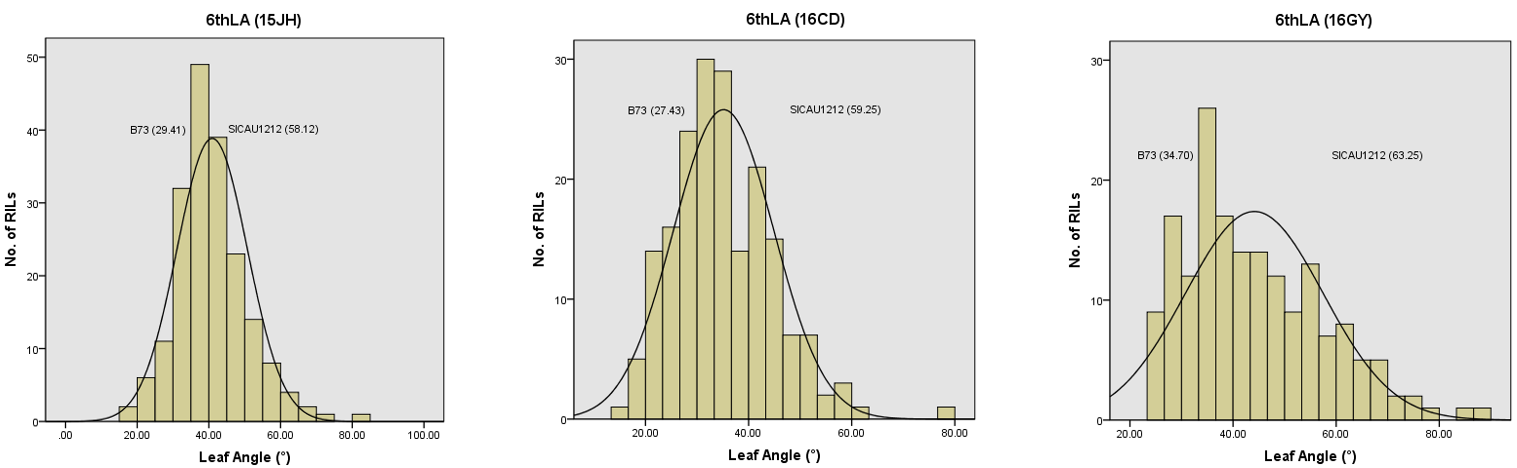

**Figure S1 continued**

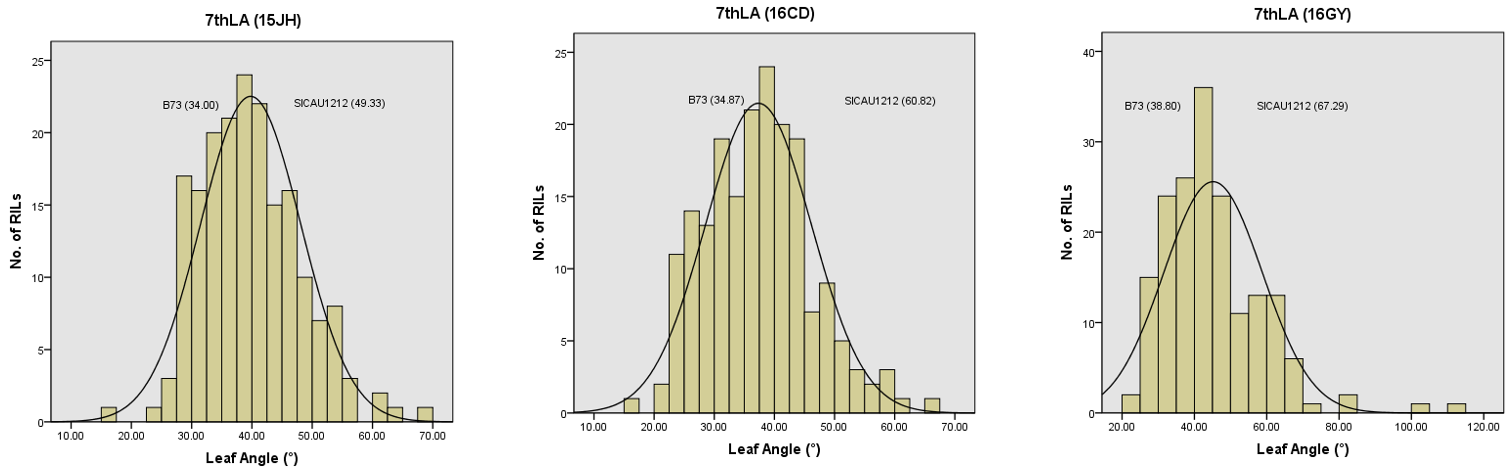

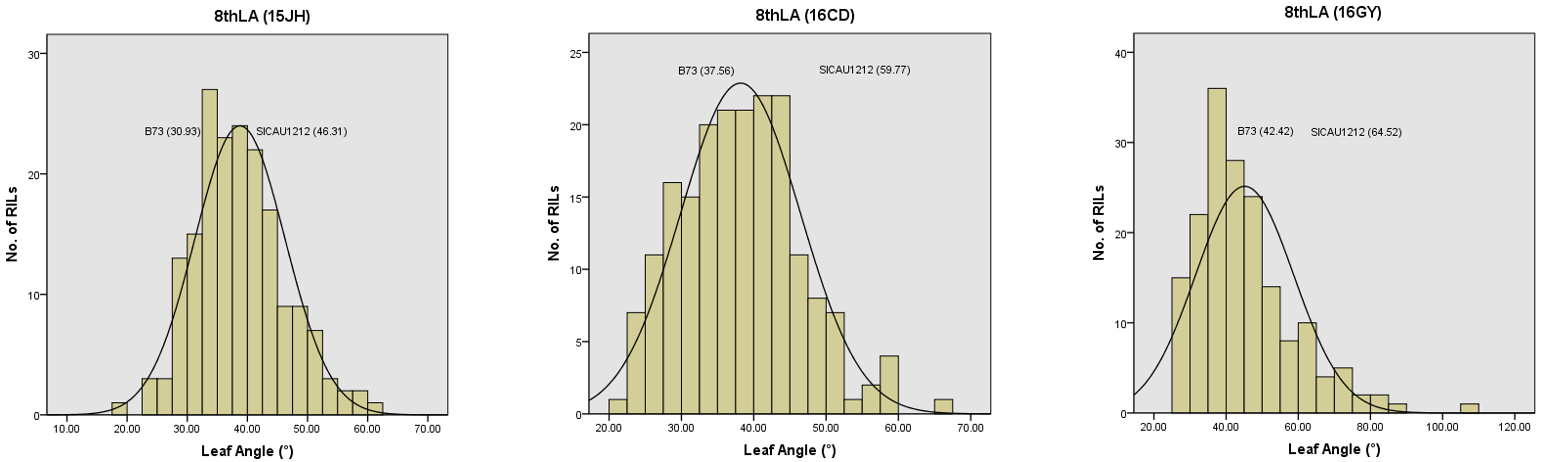

**Figure S1** Frequency distribution of LA from eight nodes in the RIL population under three environments

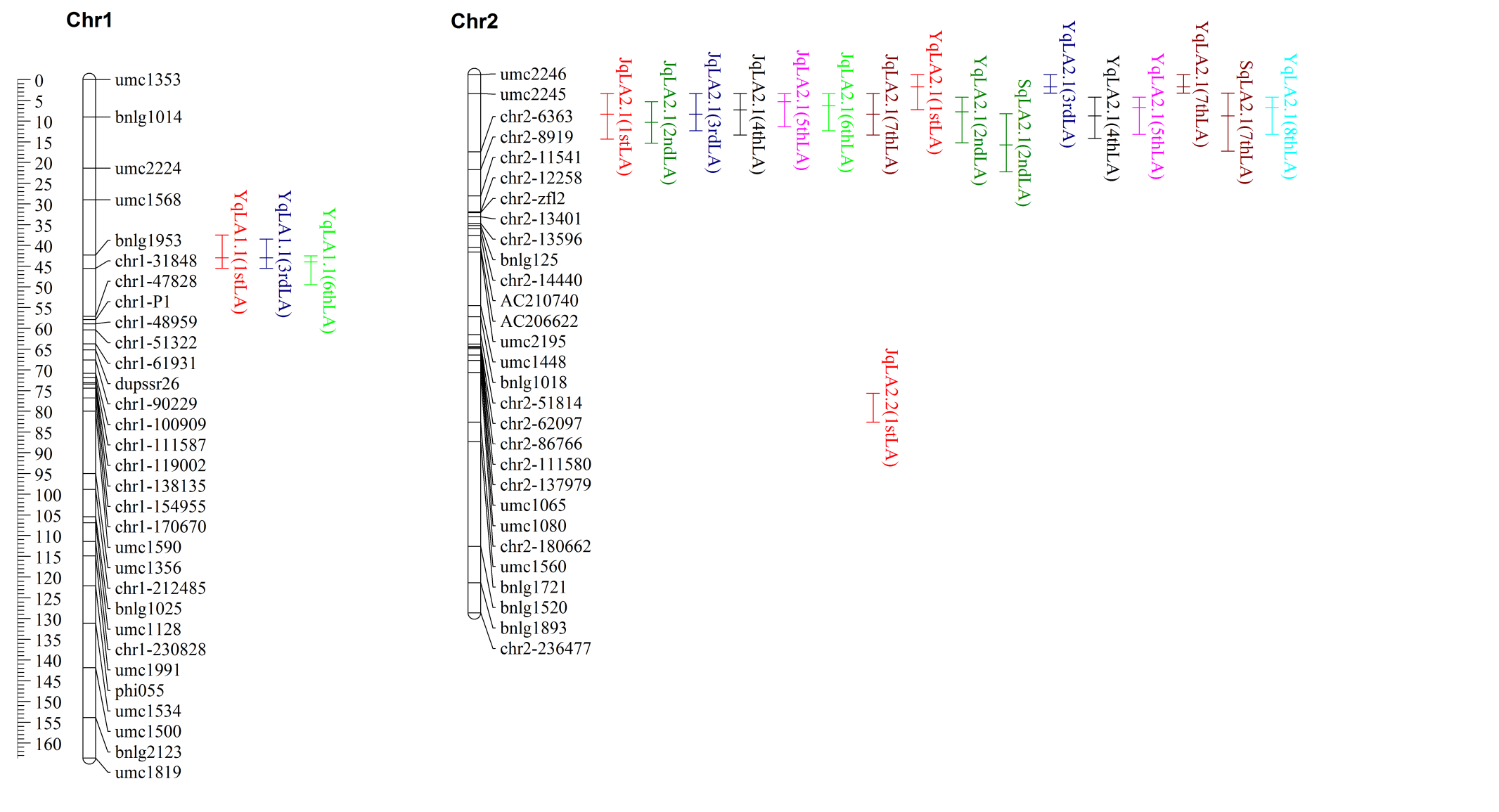

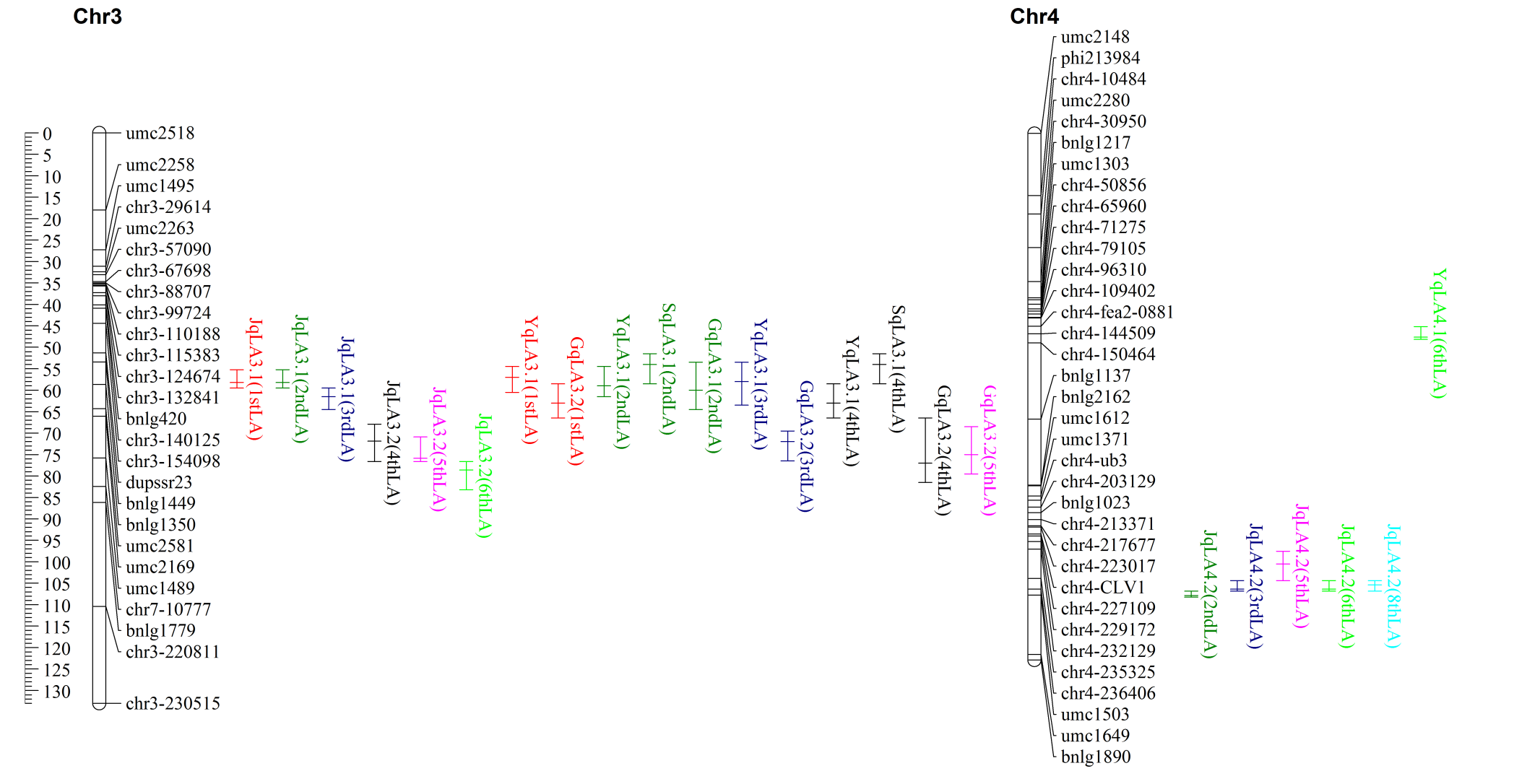

**Figure S2 continued**

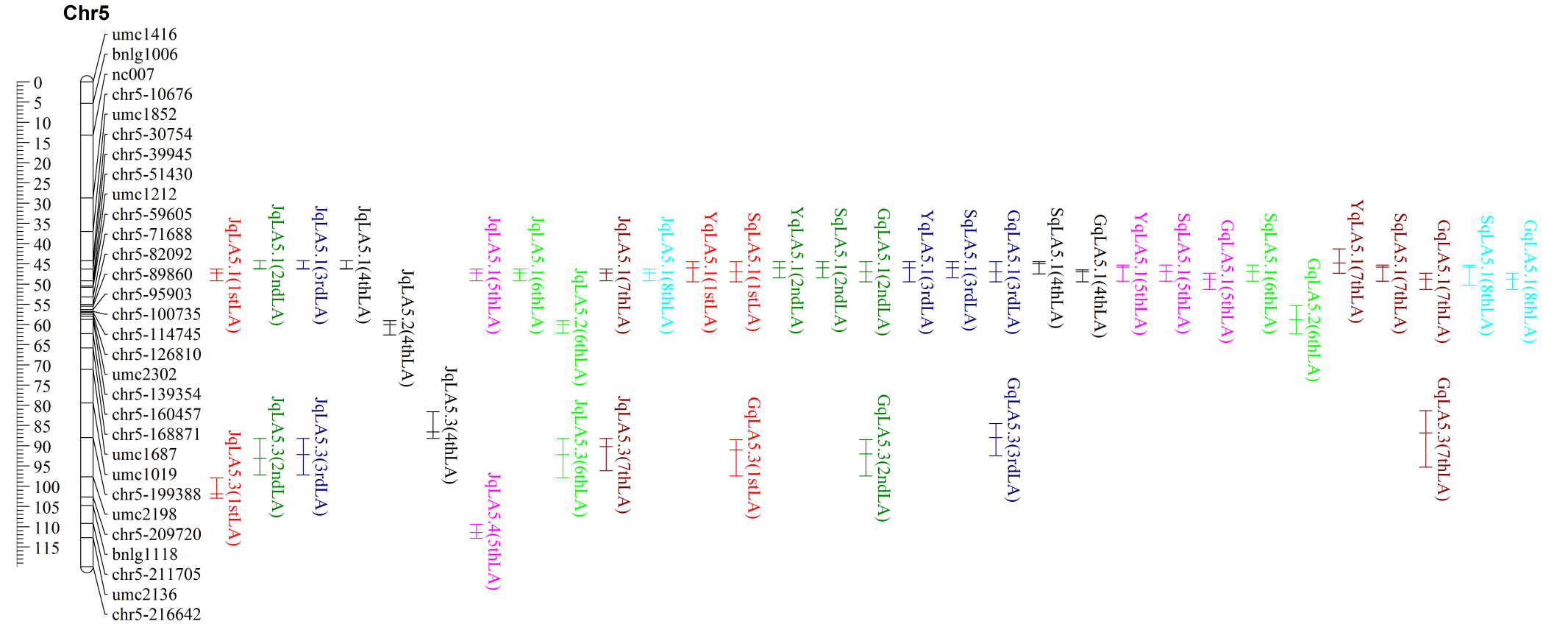

**Figure S2 continued**

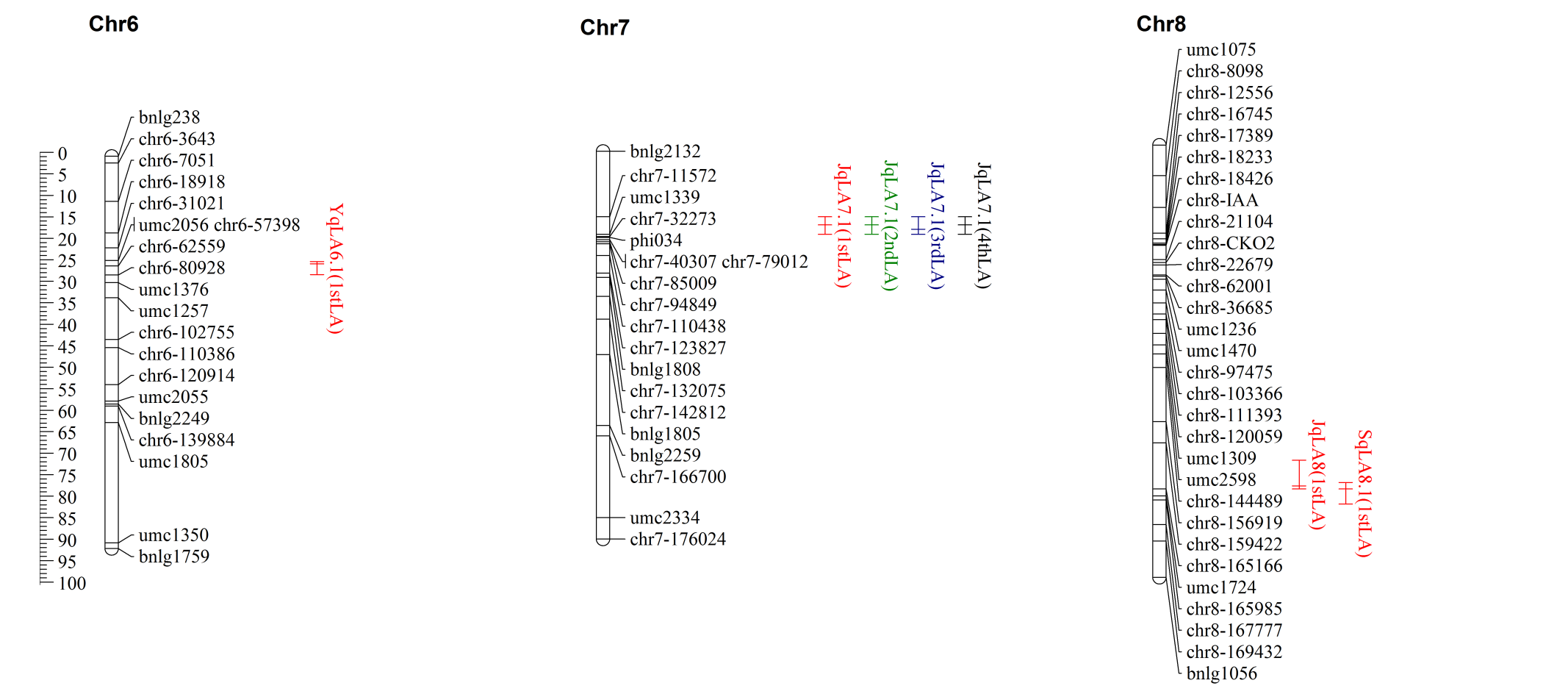

**Figure S2 continued**

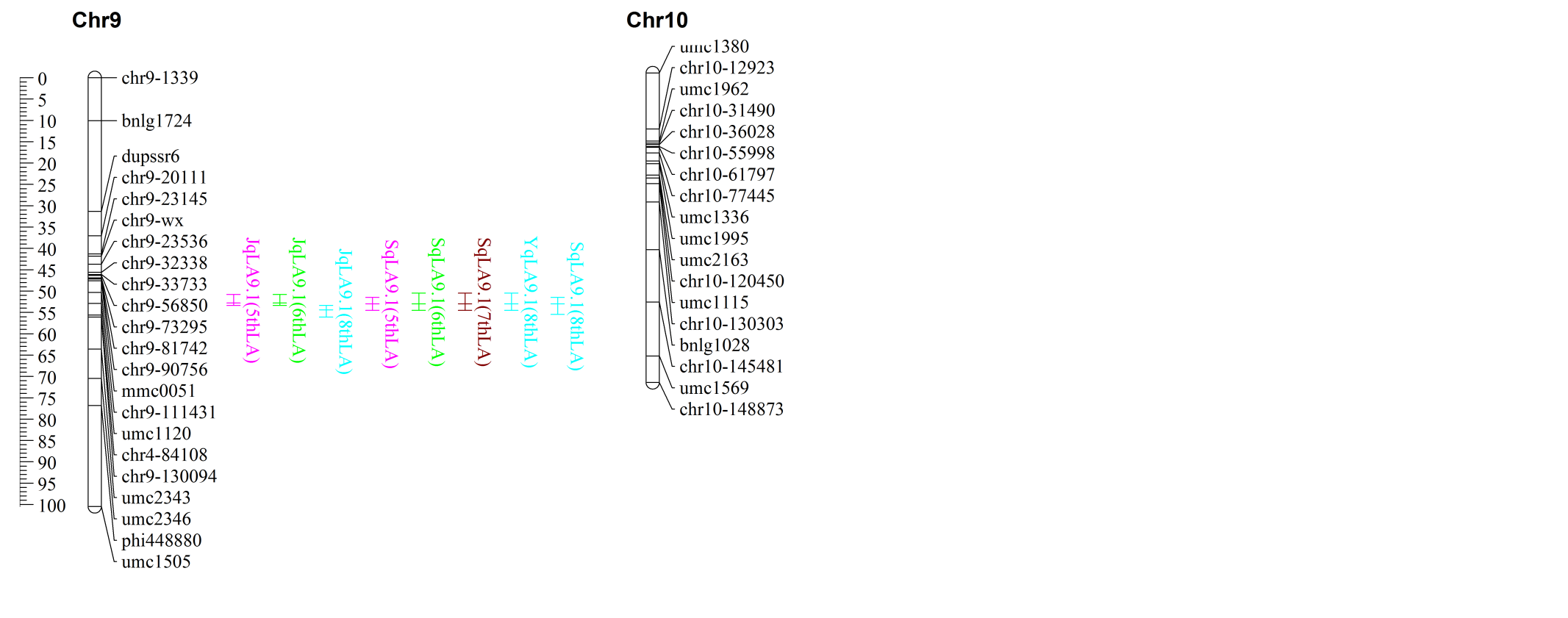

**Figure S2** Genetic linkage map of the RIL population and locations of QTL for eight LAs in single-environment QTL mapping and joint analysis. The genetic distances (cM) are indicated on the left and names of markers are shown on the right of linkage group. QTL confidence intervals were indicated by three vertical lines to the right of the linkage groups. The capital letters Y, S, G and J in the front of the name of QTL (such as YpLA1.1(1stLA)) represent QTL was detected at Jinghong of Yunnan province in 2015, Chengdu of Sichuan province in 2016, Guiyang of Guizhou province in 2016 and joint analysis across three environments, respectively.
