## Supplementary Table S1 for "Identification of QTL for leaf angle at canopy-wide levels in maize"

**Table S1.** Phenotypic correlation coefficients between eight LAs in three environments

| Traits | Env. | 1stLA | 2ndLA | 3rdLA | 4thLA | 5thLA | 6thLA | 7thLA | 8thLA |
| --- | --- | --- | --- | --- | --- | --- | --- | --- | --- |
| 1stLA | 15JH | 1 |  |  |  |  |  |  |  |
|  | 16CD | 1 |  |  |  |  |  |  |  |
|  | 16GY | 1 |  |  |  |  |  |  |  |
| 2ndLA | 15JH | 0.905^***^ | 1 |  |  |  |  |  |  |
|  | 16CD | 0.788^***^ | 1 |  |  |  |  |  |  |
|  | 16GY | 0.887^***^ | 1 |  |  |  |  |  |  |
| 3rdLA | 15JH | 0.826^***^ | 0.907^***^ | 1 |  |  |  |  |  |
|  | 16CD | 0.741^***^ | 0.872^***^ | 1 |  |  |  |  |  |
|  | 16GY | 0.862^***^ | 0.940^***^ | 1 |  |  |  |  |  |
| 4thLA | 15JH | 0.739^***^ | 0.837^***^ | 0.858^***^ | 1 |  |  |  |  |
|  | 16CD | 0.709^***^ | 0.770^***^ | 0.864^***^ | 1 |  |  |  |  |
|  | 16GY | 0.806^***^ | 0.888^***^ | 0.948^***^ | 1 |  |  |  |  |
| 5thLA | 15JH | 0.695^***^ | 0.769^***^ | 0.842^***^ | 0.849^***^ | 1 |  |  |  |
|  | 16CD | 0.656^***^ | 0.695^***^ | 0.812^***^ | 0.859^***^ | 1 |  |  |  |
|  | 16GY | 0.726^***^ | 0.797^***^ | 0.864^***^ | 0.896^***^ | 1 |  |  |  |
| 6thLA | 15JH | 0.664^***^ | 0.714^***^ | 0.769^***^ | 0.775^***^ | 0.891^***^ | 1 |  |  |
|  | 16CD | 0.619^***^ | 0.611^***^ | 0.705^***^ | 0.781^***^ | 0.898^***^ | 1 |  |  |
|  | 16GY | 0.681^***^ | 0.746^***^ | 0.769^***^ | 0.811^***^ | 0.880^***^ | 1 |  |  |
| 7thLA | 15JH | 0.563^***^ | 0.618^***^ | 0.640^***^ | 0.658^***^ | 0.777^***^ | 0.823^***^ | 1 |  |
|  | 16CD | 0.541^***^ | 0.550^***^ | 0.659^***^ | 0.731^***^ | 0.811^***^ | 0.860^***^ | 1 |  |
|  | 16GY | 0.635^***^ | 0.719^***^ | 0.743^***^ | 0.754^***^ | 0.789^***^ | 0.869^***^ | 1 |  |
| 8thLA | 15JH | 0.502^***^ | 0.577^**v^ | 0.610^***^ | 0.653^***^ | 0.775^***^ | 0.816^***^ | 0.888^***^ | 1 |
|  | 16CD | 0.453^***^ | 0.449^***^ | 0.558^***^ | 0.632^***^ | 0.753^***^ | 0.796^***^ | 0.879^***^ | 1 |
|  | 16GY | 0.606^***^ | 0.664^***^ | 0.684^***^ | 0.704^***^ | 0.726^***^ | 0.763^***^ | 0.822^***^ | 1 |

15JH, 16CD and 16GY represent Jinghong of Yunnan province in 2015, Chengdu of Sichuan province and Guiyang of Guizhou province in 2016, respectively

*** indicates significant level at *P* < 0.001
